# Somites are a source of nephron progenitors in zebrafish

**DOI:** 10.1101/2025.04.15.649022

**Authors:** Zhenzhen Peng, Thitinee Vanichapol, Phong Dang Nguyen, Hao-Han George Chang, Katrinka M. Kocha, Lori L. O’Brien, Peter D. Currie, Peng Huang, Alan J. Davidson

## Abstract

For over a century it has been believed that the vertebrate kidney arises exclusively from the intermediate mesoderm. Here, we overturn this paradigm by demonstrating that some nephrons, the functional units of the kidney, originate from the somites--blocks of paraxial mesoderm best known for their contribution to muscle and connective tissues. Using a combination of the GESTALT technique to assign developmental ancestry, somite transplantation experiments and Cre-lox fate-mapping, we show that somites can contribute to the nephrons in the adult zebrafish kidney. Our findings uncover an unexpected developmental connection between the somites and kidneys, potentially offering new pathways for developing regenerative treatments for kidney diseases.

---

Classical embryological studies in the early 20^th^ century have shaped our understanding of vertebrate kidney development, with textbooks consistently portraying the intermediate mesoderm (IM) as the exclusive source of renal progenitors^[1–3]^. It has been believed that the IM, positioned between the paraxial mesoderm and lateral plate, gives rise to a succession of different kidney types starting with the pronephros, then the mesonephros, and finally, in amniotes like birds and mammals, the metanephros. This has led to the widely accepted model that all kidney types, regardless of the vertebrate species, descend sequentially from the IM and within a relatively narrow developmental time window. However, zebrafish kidney development challenges this model. Compared to the mouse kidney development, the zebrafish pronephros forms early from the entirety of the IM, while the mesonephros develops after a significant temporal gap from nephron progenitor cells (NPCs) whose origin is unclear. The relationship of these NPCs to the IM, which appears to have long since differentiated into the pronephros, presents a conundrum with regard to the current paradigm, and calls into question the lineage relationship between the IM and the mesonephros in teleosts. Indeed, a careful histological analysis of mesonephros formation in Sandkhol Carp larvae suggests that presumptive NPCs first originate near the base of the somites—blocks of mesodermal progenitors best known for their contributions to skeletal muscle, dermis, and vertebrae^[4–7]^. Consistent with this, we found that zebrafish NPCs, which are fluorescently labelled in the Tg(*lhx1a:EGFP*) transgenic line^[8, 9]^, first appear at the ventromedial border of the anteriormost somites, adjacent to the pronephric tubules (Figure 1a). From here they migrate and expand in number around the axial vessels (Extended Data Figure 1a,b)^[8]^.

**Figure 1.**
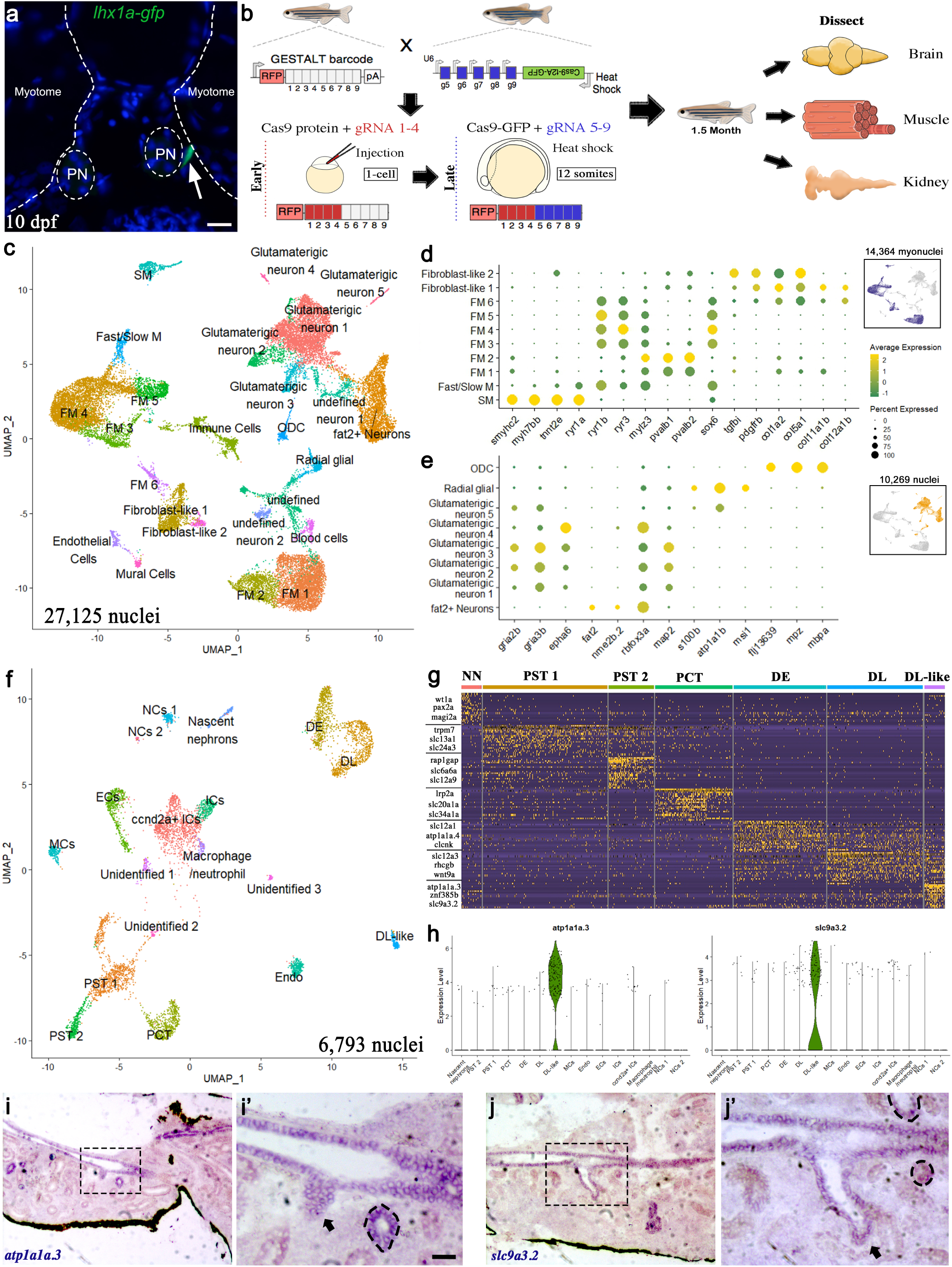
snRNA-Seq identifies various cell types from zebrafish muscle, kidney, and brain. **(a)** cross-section of 10 dpf zebrafish larvae showing the *lhx1a*-GFP^+^ kidney progenitor cells located at the interstitial region between the myotome and pronephros (scale bar, 20µm). **(b)** Schematics showing GESTALT barcode editing and tissue harvest protocol for transcriptomes and lineage recordings analysis. **(c)** UMAP of 27,125 nuclei clustered into muscle and brain cell types (n=4 independent animals). **(d,e)** dotplot of gene expression patterns of select marker genes (rows) for identified subcluster (columns) from muscle (n=14,364 myonuclei) and brain (n=10,269 nuclei). Dot size represents the percentage of nuclei expressing the marker; color represents the average scaled expression level. Inset highlights (blue and orange) the muscle and brain clusters in **(c)**. **(f)** UMAP of 6,793 nuclei clustered into kidney cell types (n=3 independent animals). **(g)** Heat map of scaled gene expression of top 20 differentially expressed genes from each nephron segment (Supplementary table S4). Key marker genes for each nephron segment are indicated on the left. NN: nascent nephrons; PST1: proximal straight tubule 1; PST2: proximal straight tubule 2; PCT: Proximal convoluted tubule; DE: Distal early tubule; DL: Distal late tubule. **(h)** Violin plot of specific markers of the DL-like tubular population. **(i)** Expression of *atp1a1a*.3 in kidney cross-sections (scale bar, 100µm). **(i’)**, close-up image of black-dashed box in **(i)**, black-dotted circle demarcates mesonephric tubule expressing *atp1a1a.3* (scale bar, 20µm). **(j)** Expression of *slc9a3.2* in kidney cross-sections. **(j’)**, close-up image of black-dashed box in **(j**), black-dotted circles demarcate mesonephric tubule expressing *slc9a3.2*.

To explore the hypothesis that these NPCs originate from the somites, we performed a lineage mapping analysis using a modified version of the scGESTALT (single cell transcriptome and genome editing of synthetic target arrays for lineage tracing) technique^[10]^. This approach maps developmental relationships between different lineages of cells using a combination of genetic barcode editing by CRISPR/Cas9 and single cell RNA-Sequencing. Transgenic zebrafish carrying the heat shock inducible GESTALT barcode (sites 1-9) were crossed to a heat shock-inducible Cas9 line (constitutively expressing guide RNAs to sites 5-9) and injected at the single-cell stage with Cas9 protein and guide RNAs 1-4, which will induce early editing at barcode sites 1-4 (Figure 1b). The embryos were then heat shocked at the 12-somite stage to induce a second round of editing at sites 5-9 (Figure 1b). Larvae with successful editing at both early and late barcode sites (Extended Data Figure 1c) were raised to 1.5 months of age (∼1 cm in length), the kidney and muscle were isolated, together with the brain as a control. Rather than performing scRNA-Seq, we chose instead to undertake single nuclei RNA-Seq, due to the multinuclear nature of some muscle fibres (herein we refer to the analysis as single nuclear (sn) GESTALT). Therefore, the nuclei were purified from each tissue, the brain and muscle samples were pooled together, and the nuclei were sorted and encapsulated using the 10X Chromium platform. We recovered the transcriptomes from four brain/muscle samples and three kidney samples. Using the unsupervised modularity-based clustering approach (UMAP), nuclei were grouped into clusters. 31 transcriptionally distinct muscle and brain populations (Extended Data Figure 1d,e) and 18 transcriptionally-distinct kidney populations (Extended Data Figure 1i) were resolved. Established anchor markers reported in the literature and the ZFIN database were then used to assign presumptive cell identities to each population. Clusters expressing *Titin* genes (*ttn.1*, *ttn.2*) and myosin heavy chain genes (*myhc4*) were classified as muscle nuclei (n=14,364), and clusters expressing neuronal genes (*luzp2*, *nrxn3a* and *grid1b*) were classified as brain nuclei (n=10,269) (Extended Data Figure 1f). This resulted in the identification of 12 muscle and 9 brain populations. In addition, we found presumptive populations of immune (*ptprc*), endothelial (*egfl7*), blood (*hbaa1*) and mural cells (*myh11a*) (Extended Data Figure 1h). Three populations (cluster 12, 19 and 24) could not be assigned to either muscle or brain, due to a lack of expression of anchor genes.

To further classify the muscle clusters, we compared the differentially expressed genes in each cluster with markers specific to muscle cell types (fast and slow), as well as developmental stage (Supplementary table 1)^[11–13]^. From this we identified 6 distinct fast muscle (FM) subtypes (FM 1-6 in Figure 1c,d). FM1 and FM2 show enriched expression of the parvalbumin genes, *pvalb1* and *pvalb2*, which play a role in fish muscle relaxation^[14]^. FM3 to FM6 show high expression of the ryanodine receptor genes, *ryr1b* and *ryr3*, *mef2cb*, *msi2b* and *sox6*, which are markers of developing fast muscle fibers (Figure 1d and Extended Data Figure 1g) ^[15]^. Given that FM1 and FM2 express lower levels of the pro-differentiation genes, we speculate that these clusters correspond to more mature fast muscle fibers cells. We identified one slow muscle fiber population (SM) that expresses markers such as *smyhc2*, *myh7bb*, *mybpc1* and *tnnt2e* ^[11]^ (Figure 1d). In addition, we found a mixed fast/slow population that expresses *ryr1b* and *ryr3,* but also the slow muscle fiber gene *ryr1a*, suggestive of common progenitor population of intermediate fibres^[6]^ (Figure 1d). Two mixed fibroblast populations that expressed several collagen genes were also identified (Figure 1d).

For the brain nuclei, we grouped them into 9 transcriptionally-distinct neuron populations that included 5 glutamatergic subtypes (expressing *gria2b*, *gria3b* and *epha6*), one *fat2^+^* subtype and 2 undefined subtypes (Figure 1c,e and Supplementary table 2). All of these populations, except for glutamatergic neuron-5, are likely mature neurons, as they expressed the markers *rbfox3a* and *map2* ^[16, 17]^. Our analysis also identified a presumptive group of neuronal stem cells, radial glia cells (expressing *s100b*, *atp1a1b* and *msi1*), and oligodendrocytes (expressing *flj13639*, *mpz* and *mbpa*; Figure 1c,e).

Nuclei from the kidney were grouped into populations corresponding to the nephron segments (7 subtypes, n=3,854 nuclei), haematopoietic cell lineages (6 subtypes, n=2,939 nuclei) and neurons (2 subtypes, n=214 nuclei) (Figure 1f, Extended Data Figure 1j,k). These clusters were annotated with anchor genes, as well as differentially expressed genes^[8, 18, 19]^ (Supplementary table 3). While no podocytes or NPC clusters were identified, we did resolve proximal and distal tubule nephron populations. These comprised one proximal convoluted tubule population (PCT), which was defined by expression of *lrp2a*, *slc20a1a* and *slc34a1a*^[20–22]^, and two proximal straight tubule populations (PST1 and PST2), based on the expression of *trpm7* and *slc13a1*^[21–23]^ (Figure 1g). Differential gene expression analysis between these subtypes revealed that PST2 also expresses *rap1gap*, *slc6a6a* and *slc12a9*. For the distal nephron segments, we identified distal early (DE) cells, which express *clcnk* and *slc12a1*, and distal late (DL) cells that express *wnt9a* and *slc12a3*^[22–24]^. We also have identified a new DL-like population that specifically expresses *atp1a1a.3* and *scl9a3.2* (Figure 1h). GO Term analysis for this population is consistent with a role in ion transport (Extended Data Figure 1l). To determine the spatial location of this DL-like segment we performed *in situ* hybridisation on adult kidney sections and found expression in the major collecting ducts (running down the midline of the kidney) and in the distal-most portion of nephrons that fuse with the collecting ducts (Figure 1i,j).

To determine the relationship of cells from kidney and muscle using snGESTALT, we prepared and sequenced the snGESTALT lineage libraries, resulting in barcode recovery of 358 nuclei from two juvenile zebrafish, respectively, corresponding to ∼2% of all profiled cells. We filtered out the barcodes with only one UMI (Unique Molecular Identifiers) and constructed lineage trees for the remaining barcodes using the published maximum parsimony approach^[10]^, anchoring the tree with edits at sites 1-4 and extended it with edits at sites 5-9. Although the recovery rate of the barcodes was low, most likely due to weak expression of the heat-shock promoter and lower recovery from nuclei, they cover the majority of the identified populations in all three tissues (Extended Data Figure 2a,b). A shared barcode between different types of cells indicates a shared ancestry and we found that the majority of these were restricted to single tissue types e.g. between different types of muscle cells (branch a3, b2, Figure 2A; branch a1, a2 and b2. Figure 2b) or between nephron segments (branch a1, b1, Figure 2a) or different neuronal cells (branch b3, Extended Data Figure 2a). We found one barcode shared by a fast muscle cell (FM5) and a *fat2*^+^ neuronal cell, likely reflecting the lineage relationship between bipotent neural mesodermal progenitors that contribute to the spinal cord and somitic mesoderm^[25]^ (branch a3 in Extended Data Figure 2b). No shared barcodes were found between the kidney and brain (Extended Data Figure 2). However, we found two examples of barcode sharing between the muscle and kidney lineages. The first of these revealed a lineage relationship between an FM3 cell and a DL-like tubular cell (Fish 1 branch a2, Figure 2a), while the other linked an FM4 cell to three tubular cells (PST1, PST2 and DL, Fish 2 branch b1, Figure 2b). These findings are consistent with the hypothesis that the somites can give rise to kidney tissue.

**Figure 2.**
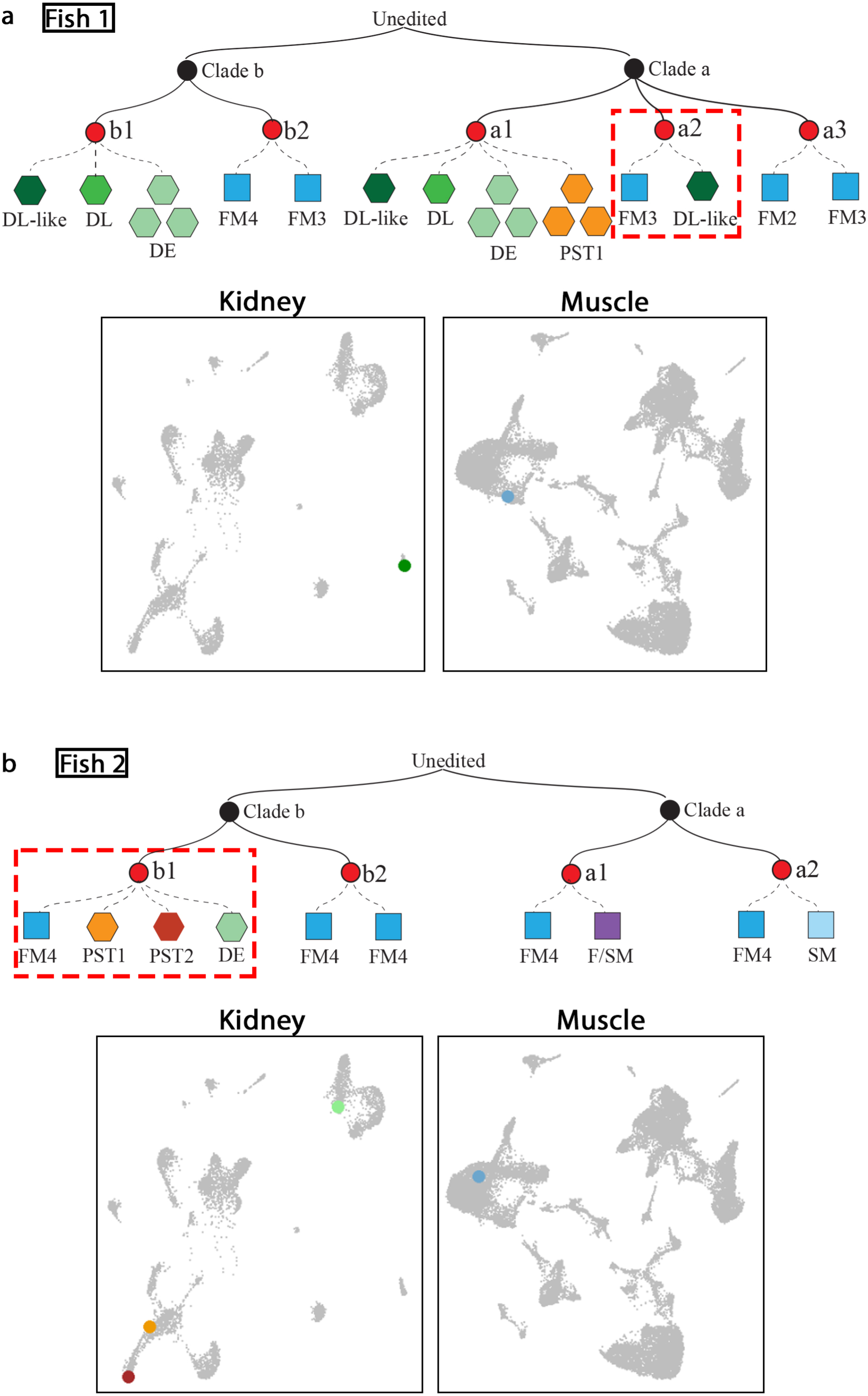
Reconstructed lineage trees reveal linkage between muscle and kidney cells in juvenile zebrafish. **(a)** Mini-tree showing lineage branches of kidney and muscle cells from fish 1 sharing the same barcode (example branches from Figure S2A). **(b)** Mini-tree showing lineage branches of kidney and muscle cells from fish sharing the same barcode (example branches from Figure S2B). Black nodes indicate early edits, red nodes, late edits. Black solid lines represent lineage relationships between nodes. Dashed lines connect individual cells to the nodes and represent the cells that share the same barcode. Each shape represents a cell coloured by cell type. Right, UMAP plots with highlighted cells from kidney and muscle share the same barcodes, red dash boxed highlighted branches.

To further test whether somites are a source of NPCs, we performed somite transplantation experiments. We selected the 3rd somite, based on its proximity to the where the first NPCs will arise later development, and dissected this from transgenic embryos ubiquitously expressing *egfp* at either the 12-or 18-somite stages. These donor somites were then transplanted into equivalently staged unlabelled recipients where the corresponding somite was removed (Figure 3a and see Extended Data Figure 3a-e for an overview of the transplant procedure). In surviving recipients, we observed successful engraftment through the presence of GFP^+^ donor-derived muscle fibres at the transplantation site (Figure 3b). Follow-up analysis at 1.5 months revealed continued donor-derived contribution to muscle fibres at the transplantation site as well as donor-derived Hnf1b^+^ mesonephric kidney tubules and glomeruli in the mesonephros (n=5/100 recipients, with n=3 with 12-somite stage donors and n=2 from 18-somite stage donors; Figure 3c-d and Extended Data Figure 3f-h). Notably, in one recipient, donor-derived tubules were found contralateral to the transplantation side, consistent with our previous observations that NPCs exhibit migratory behaviour^[18]^ (Figure 3c’).

**Figure 3.**
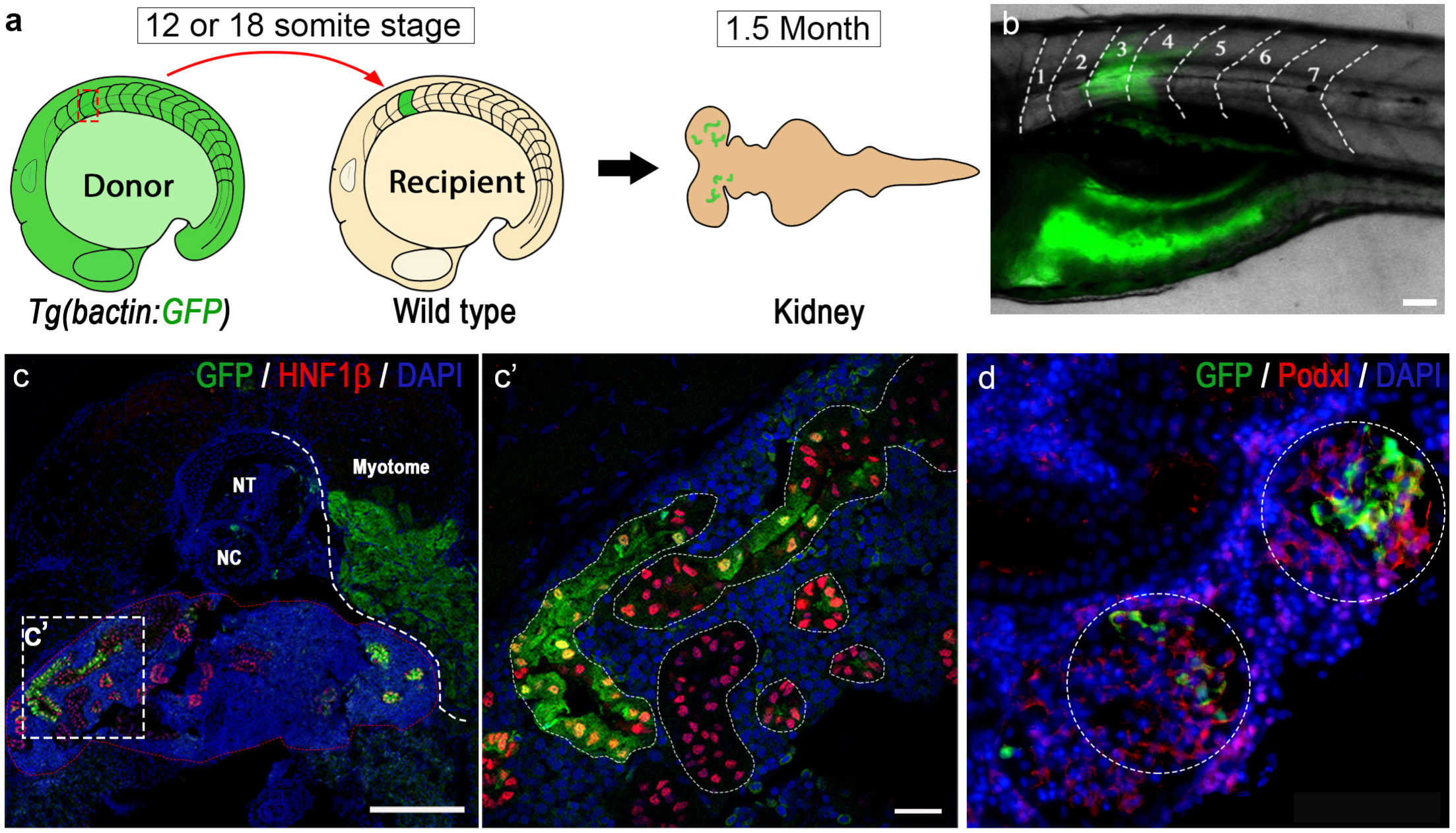
Transplanted GFP^+^ somite contributed to mesonephric nephrons in recipient fish. **(a)** Schematic of somite transplantation showing a GFP^+^ donor somite is harvested at 12 or 18 somite stage and transplanted into a wild type recipient embryo at equivalent developmental stage, and kidney of the recipient fish is examined at 1.5-month of the age. **(b)** Image of a recipient larvae (6 dpf) developed donor derived GFP^+^ muscle fibers at the transplantation site (somite 3, transplanted at 12 somite stage). Scale bar, 20µm. **(c)** Cross section of same recipient juvenile fish (at 1.5-month stage) immunostained with GFP and renal tubular marker Hnf1b, showing donor somite contributed tubular structures (Scale bar, 100µm). **(c’),** close-up image of the area boxed in c showing Hnf1b^+^ mesonephric tubules containing donor somite derived GFP^+^ cells (Scale bar, 20µm). **(d)** Immunostaining of donor derived Podxl^+^/GFP^+^ glomeruli, n=3, Scale bar, 20µm.

To provide further evidence supporting a renal-somite lineage relationship, we conducted conventional 4-hydroxytamoxifen (4-OHT)-inducible Cre-mediated (CreERT2) genetic fate-mapping experiments *in vivo* using transgenic lines employing the paraxial mesoderm-specific *mesogenin1* (*msgn1*) ^[26]^ and the sclerotome marker *NK3 homeobox 1* (*nkx3.1)* promoters^[27]^ (Figure 4a). In *msgn1*:*Cre-ERT2*; *βactin:Switch* fish, paraxial mesoderm-derived cells were induced to express mCherry with 4-OHT treatment at the 50% epiboly stage (5.25 hpf), while in *nkx3.1:Gal4; UAS:Cre-ERT2; ubi:Switch* fish, sclerotome-derived cells were switched at 24 hpf (Figure 4b). Both lines were treated with 4-OHT for a duration of 24 hours. As expected, the mCherry^+^ cells derived from the *msgn1* lineage gave rise to muscle fibres at 48 hpf (Extended Data Figure 4b,c), while no labelling of the IM-derived pronephros was observed at either 48 hpf or at 2 weeks post-fertilisation (7 mm stage; Extended Data Figure 4d and Figure 4c and 4d, n=5). The mCherry^+^ cells derived from the *nkx3.1* lineage were found associated with the myotendinous junction, centrum, neural arch and hemal arch (as reported by Ma *et al*. 2023, Extended Data Figure 4f). Consistently, some mCherry^+^ muscle fibres were also observed in the ventral part of myotome (Extended Data Figure 4f, yellow arrow heads). Embryos of both lines without 4-OHT induced Cre showed no presence of mCherry^+^ cells (Extended Data Figure 4e,f). We examined the recombined animals from both lines at the 7 mm stage (∼3 weeks post-fertilization), a crucial period when the mesonephric kidney begins to form, and we observed contribution of mCherry^+^ cells to nascent Pax2^+^ nephrons (n=5; Figure 4c) and NPC clusters (n=5; Figure 4e). Additionally, we observed contribution of mCherry^+^ cells to Hnf1b^+^ nephrons in both lines (n=10; Figure 4d and f), including mCherry^+^ glomeruli (Figure 4f, n=10). Recombined animals (1.5 months of age) of both lines exhibited mCherry labelling in nephrons scattered throughout the kidney with mosaic contribution in the glomeruli, proximal and distal tubules (Figure 4g, 4h and 4i, n=10 per line). Additionally, 4-OHT was administered to *msgn1* embryos at the later stages of 3-and 10-somites, where there is no overlap in expression between *msgn1* and the IM marker *pax2a* (Extended Data Figure 4g). A similar efficiency of nephron contribution was found among the three different 4-OHT treatment stages (Figure 4o), although the labeling of muscle fibers was more restricted to the posterior somites in the 10-somite treated animals compared to the earlier treatment windows (Extended Data Figure 4h, Figure 4m). The *nkx3.1* line demonstrated 6-fold greater efficiency in contributing to nephron formation compared to *msgn1* line (Figure 4n and 4o). Taken together, the data from both *msgn1* and *nkx3.1* studies provide further independent evidence for a somite origin of NPCs.

**Figure 4.**
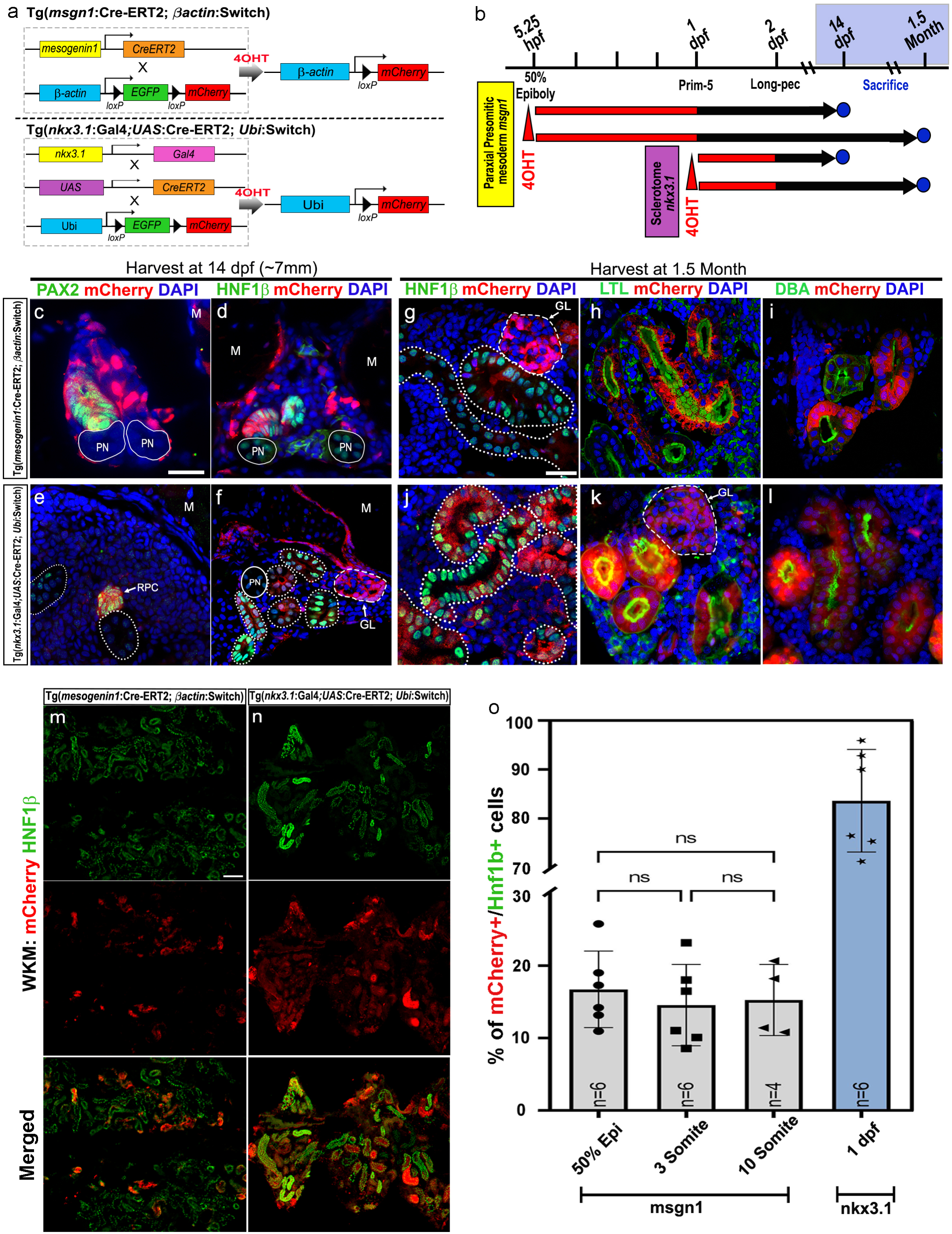
Cre-loxP based lineage-tracing techniques unveil a somitic origin for zebrafish mesonephric nephrons. **(a)** Schematic representation showing the cross between the *msgn1*:Cre-ERT2 line and *bactin*:Switch (loxP-EGFP-loxP-mCherry) line, and the *nkx3.1*:Gal4;UAS:Cre-ERT2 line and *Ubi*:Switch line, followed by treatment with 4-OHT to activate Cre recombinase and induce mCherry expression in targeting cells. **(b)** Experimental timeline of 4-OHT treatment in both transgenic lines. **(c-f)** Cross-sections of 4-OHT treated 7mm larvae immunostained for renal markers (Pax2, Hnf1β) and mCherry^+^ cells. *Mesogenin1*-Cre X *bactin*-Switch **(c-d)** and *nkx3.1*-Gal-UAS-Cre X *Ubi*-Switch **(e-f)**, (Scale bar, 20µm); M: muscle, RPC: renal progenitor clusters. **(g-l)** Sections of 4-OHT treated 1.5-month zebrafish kidney immunostained for renal markers (HNF1β, LTL, and DBA) and mCherry^+^ cells, showing their contribution to glomerulus (GL, white dash outlines), proximal tubule (LTL), and distal tubule (DBA) of the nephron (white dotted outlines). PN: pronephros (solid white outlines), Scale bar, 20µm. **(m-n)** Representative low-magnification images of kidney sections from both transgenic lines for quantification. Scale bar, 100 µm. **(o)** Quantification of mCherry^+^ cells in 1.5-month *msgn1*-Cre and *nkx3.1*-Gal-UAS-Cre fish kidneys. The quantification of *msgn1*-Cre line was performed at three different recombination time points (50% epiboly, 3-somite, and 10-somite, n=6 fish per time point). Each dataset consists of three sections quantified per fish, with each section 14μm in thickness, and every fifth section was quantified.

In summary, we demonstrate through three independent lineage tracing methods that somites serve as a source of NPCs for mesonephric nephrons in zebrafish. This finding challenges the prevailing assumption that the IM is the exclusive origin of all kidney cell types in vertebrate embryos. This long-standing belief primarily emerged from studies of chick and mammalian embryogenesis, where the IM develops progressively as the embryonic axis elongates and forms ‘nephrogenic cords’ alongside the somites. These IM cells are the source of the sequentially-arising kidney types— the pronephros and mesonephros in the rostral trunk and the metanephros in the caudal trunk of mammalian and chick embryos^[3, 28, 29]^. By contrast, zebrafish kidney development follows a markedly different trajectory. The IM forms in its entirety by early somitogenesis stages and then differentiates *in toto* into the pronephros, with no obvious ‘remnant’ nephrogenic cord cells remaining to serve as a source for the mesonephros. Unlike the other models, the zebrafish mesonephros emerges much later in development, in more mature, free-swimming larva. The absence of persisting nephrogenic cord cells in zebrafish has long been a developmental mystery that our work now resolves by revealing that the somites are an alternative source of NPCs.

During zebrafish somitogenesis, the somite becomes compartmentalised into discrete domains that include the primary myotome (contributing to fast muscle fibres), a dermomyotome-like external cell layer, and dorsal and ventral sclerotome populations that are a source of multiple types of mesenchymal cells and the vertebrae^[27, 30, 31]^. Our lineage tracing with *msgn1*:*Cre-ERT2* does not resolve which of these compartments is the source of NPCs, as *msgn* is expressed from early gastrulation onwards throughout the paraxial mesoderm^[32]^. By contrast, *nkx3.1* is more restricted in the somite and has a later onset of expression, where it predominantly labels sclerotome progenitors^[27, 33]^. This result, coupled to the observations that the first NPCs arise near the base of the myotome, suggest that ventral sclerotome population may be the source of NPCs. Whether a subset of sclerotome progenitors persist to ∼10 dpf, when the first NPCs arise, or whether there is an intermediary cell type/s involved in the developmental pathway from somite to NPC is not clear but can be addressed in the future with additional lineage-restricted Cre-lox tools.

Alternatively, as *nkx3.1-cre* also labels some muscle fibres^[27]^, together with our GESTALT analysis where we found that some fast muscle cells share a common ancestry with kidney cells, we cannot rule out that NPCs descend from the myotome compartment. Although the trunk muscle initially arises from the primary myotome, during the larval and adult stages it is greatly expanded by the differentiation of muscle stem cells on the myotome surface^[6]^. It is possible that these muscle stem cells are the source of NPCs, and this would fit with the observations that NPCs arise during larval stages when secondary myogenesis is active, together with their first appearance at the surface of the myotome. In line with this, the GESTALT results found that the muscle cells sharing a common ancestry with the renal cells have an immature transcriptional profile, suggestive of being newly-derived during secondary myogenesis^[6]^. A link between muscle stem cells and the kidney is attractive as both tissues show lifelong indeterminant hyperplasia in teleosts^[6, 8]^ and a common stem cell population may help coordinate this growth.

Our findings raise the question of whether the somite-to-kidney developmental pathway is a unique adaptation found in teleosts or whether it reflects an ancient vertebrate developmental mechanism that has been overlooked in mammals and birds. Lineage labelling studies in chick embryos have shown that the paraxial mesoderm contributes to the mesonephros and metanephros but is restricted to stromal cells^[34]^. While the stromal and NPC populations were initially considered to be developmentally distinct lineages, more recent observations suggest a much closer relationship. Lineage tracing of mouse stromal precursors using *Foxd1-cre*, which shows earlier expression in the somites, labels the metanephric stroma but also small populations of metanephric NPCs^[35]^. Moreover, knockout of the *Pax2* transcription factor in metanephric nephron progenitors causes them to transdifferentiate into stromal progenitors^[36]^. In the developing human metanephric kidney, the transcriptional profiles of nephron and stromal progenitors overlap substantially, including shared expression of *FOXD1*, considered a specific stromal marker in mouse^[37]^. Together, these observations raise a provocative hypothesis that the somite-to-kidney pathway we describe in zebrafish may also exist in mammals and birds but it has become co-opted to generate stromal, rather than nephron, progenitors.

In conclusion, our work challenges the traditional view of the IM as the sole source of NPCs. It extends the remarkable plasticity of somites as a key developmental reservoir of tissue progenitors and offers a provocative lens through which to reconsider amniote kidney development. Our research opens the possibility of inducing human nephron progenitors *in vitro* by manipulating a somite-to-kidney pathway and utilising them in regenerative therapies aimed at treating kidney disease.

## RESOURCE AVAILABILITY

### Lead contact

Further information and requests for resources should be directed to and will be fulfilled by the lead contact, Alan J Davidson

### Data and Code availability

The sequencing data for this study have been deposited in the Gene Expression Omnibus under accession number GSE286280. All snRNA-seq data analyses were performed using standard protocols with previously described R packages^[38]^. GESTALT data were processed using Single-cell GESTALT pipeline available at GitHub (https://github.com/mckennalab/SingleCellLineage). The R scripts are available upon request.

## ACKNOWLEDGMENTS

We thank Prof Alexander F Schier, the Director of the Biozentrum University of Basel, Switzerland, and Dr Bushra Raj from the Department of Molecular and Cellular Biology, Harvard University, Cambridge Massachusetts, USA, for their technical support on GESTALT experiment and data analysis. We also thank Dr. Nikki Freed and Dr. Jieyun Wu from the sequencing facility, Auckland Genomics, the University of Auckland, for technical assistance with snRNA-seq and GESTALT library sequencing.

This study was supported by grant 18-UOA-151 from the Marsden Fund Royal Society of New Zealand (A.J.D), NHMRC Australia Investigator Grant 2016338 (P.D.C), Canadian Institutes of Health Research PJT-169113 (P.H) and NIH R01DK121014 (L.L.O.).

## AUTHOR CONTRIBUTIONS

Conceptualization, Z.P. and A.J.D.; investigation, Z.P., T.V., P.D.N., and H.G.C.; visualization, Z.P., T.V., A.J.D.; snRNA-Seq analysis, Z.P. and T.V.; formal analysis, Z.P. and A.J.D.; resources, P.D.N., P.T.D., K.K., and P.H.; writing – original draft, Z.P., and A.J.D; writing – review and editing, P.H., P.D.C., L.L.O.; supervision, A.J.D; Project administration, Z.P. and A.J.D.; funding acquisition, Z.P. and A.J.D.

## Supplemental information

Supplementary Table 1. DEGs for muscle clusters.

Supplementary Table 2. DEGs for brain clusters.

Supplementary Table 3. DEGs for kidney clusters.

Supplementary Table 4. Top20 genes for each kidney cluster.

Supplementary Table 5. The sequence information for primers.

## METHODS

### Zebrafish husbandry

Zebrafish husbandry and manipulations were conducted at the facilities of the University of Auckland, Faculty of Medical and Health Sciences. This study was approved by the University of Auckland Animal Ethics Committee under approved protocol AEC22634.

### Zebrafish transgenic lines

Zebrafish were maintained as previously described^[39]^ and according to Institutional Animal Care and Use committee protocols. The transgenic lines Tg(*lhx1a:EGFP*), Tg(*hsp70l:Cas9-t2A-GFP, 5xU6:sgRNA*), Tg(*hspDRv7:GESTALT, clmc2:EGFP*), Tg*(bactin2:loxP AcGFP1-STOP loxP pA mCherry pA*), Tg(*msgn1:CreERT2*) were previously reported^[10, 26]^.

### Imaging

Epifluorescent images were taken from a Nikon Eclipse 80i microscope using the Nikon Qi2 camera and confocal images were acquired using the Zeiss LSM 710 inverted confocal microscope.

### Cryosectioning

Zebrafish larvae and kidneys were fixed in 4% PFA overnight at 4℃ and washed twice in PBS and then placed in 1% low melting point agarose containing 5% sucrose and 0.9% agar (made up in water) in cryomoulds and placed in 30% Sucrose/PBS cryoprotectant solution and incubated overnight at 4℃. Cryoblocks were flash frozen on dry ice and immediately sectioned. 14 μm sections were cut in a cryostat machine (Leica SM-3050-S) with chamber temperature set to −25℃ and objective chamber set to −22℃, and transferred onto Superfrost plus microscope slides.

### *In situ* hybridization and immunohistochemistry

Both *in situ* hybridization and immunohistochemistry were performed on cryosectioned samples.

*In situ* hybridization was performed as previously described (http://zfin.org/ZFIN/Methods/ThisseProtocol.html) with the sample section slides kept in the humid chamber to prevent drying during the reaction. Antisense probes were synthesized using T3 polymerase from cDNA synthesized from kidneys of 1.5-month-old juvenile fish. To perform antibody staining, tissue sections were rinsed twice in PBS containing 0.05% Tween20 (PBST), then incubated in PBS containing 0.5% Triton X100 for 20 min to permeabilise the sections. Sections were then blocked in PBST containing 3% BSA and 5% Goat serum for at least 1 hr.

Antibody staining was performed as described (https://zfin.atlassian.net/wiki/spaces/prot/pages/379420760/Antibody+staining+on+Sections), using antibodies anti Pax2 (1:500, Covance, PRB-276P-200), anti Hnf1b (1:500, Sigma, HPA002083-100UL), anti GFP (1:500, Novus, NOVNB600597), anti mCherry (1:500, Abcam, ab125096), anti Podxl (1:250, made in House by Dr. Hidetake Hurihara, Juntendo University). Sections were washed three times in PBST before the Donkey anti-mouse/Rabbit IgG (H+L) Highly cross-adsorbed Secondary antibodies (1:500, Thermo Fisher Scientific). Antigen retrieval was required for the anti-mCherry antibody using citrate solution (10mM citric acid, 0.05% Tween, pH6.0) in 95°C water bath for 20 minutes.

### Somite Transplantation

The transplantation method was adapted from a previously published work^[40]^, with several modifications made. In brief, individual donor somites were harvested from 12 or 18 somites stage β-actin-GFP transgenic zebrafish embryos. The dechorionated donor embryos were kept in L15 media for removal of the head and yolk by using two 29G insulin needles. The remaining trunk of the embryo was digested in 1% collagenase (Sigma) until the entire somites blocks started to detach from the embryo. The digested somites blocks were then washed by serial of L15 media supplemented with 10% FBS. Individual somite blocks were separated and selected under a fluorescent dissection microscope (Leica MZ F10) by using an eye lash and transferred to ice-cold fresh L15 media until transplantation. Dechorionated wild-type recipient embryos (12 or 18 somites), were embedded in 60mm Petri dish with 1% low-melting/normal agarose gel (50:50) and the embryos oriented that the lateral surfaces of the targeted somite regions were at the surface of the embedding gel. Approximately 2 mL of L15 media supplemented with 10% FBS was added to the petri dish to ensure the surface was fully covered and embryos would not dry out. Under a dissecting microscope, the epidermal layer of the target somite was cut open using two Micro-Needles (0.12mm in diameter, Ted Pella), the target somite was then hooked out using a blunt end eye lash. Immediately following the surgery, the designated donor somite was transplanted into the extirpated region of the recipient host. After transplantation, the embryos were left undisturbed and immersed in the L15 media (supplemented with 10% FBS) for at least 30 minutes before transferring to E3 medium. The embryos were incubated at 28°C for recovery, survival of the embryos was checked at 1 day post transplantation. The surviving embryos were allowed to grow to 1.5 months and analyzed under fluorescent microscope. Kidney tissue was harvested and analyzed by immunohistochemistry.

### Early and late GESTALT barcode editing

The editing was performed as previously described^[41]^, minor changes were made to fit with the experimental design of this project. In brief, sgRNAs specific to sites 1-4 of the GESTALT barcode were generated by in vitro transcription using EnGen^®^ sgRNA Synthesis Kit, *S. pyogenes.* Tg(*hspDRv7:GESTALT, clmc2:EGFP*) F1 transgenic adults confirmed with single copy of GESTALT barcode were crossed to heat-shock inducible Cas9 F1 transgenic adults. One-cell embryos were injected with 1.5nl of Cas9 protein (8µM EnGen^®^ Spy Cas9 NLS, NEB) and sgRNAs 1-4 (100 nl/µl) in salt solution (NEBuffer™ r3.1) with 0.05% phenol red to perform early GESTALT barcode editing. Injected embryos were first screened for GFP heart expression at 30 hpf to select the embryos with the presence of GESTALT barcode. These embryos were then heat-shocked for 45 minutes at 37°C to induce endogenous Cas9 expression, which performed GESTALT barcode editing at sites 5-9 (late editing).

### Nuclei extraction for snRNA-Seq

Single nuclei isolation from tissue was adapted from a previously published work ^[42]^, minor changes were made to fit with the size of zebrafish tissue. Nuclei were isolated with Nuclei EZ Lysis buffer (NUC-101, Sigma) supplemented with protease inhibitor (5892791001, Sigma) and RNase inhibitors (N2615, Promega; AM2696, Life Technologies). Zebrafish brain, muscle and kidney tissue were dissected and quickly rinsed with ice-cold PBS before snap-froze. Samples were minced into <2-mm pieces with 500 µL of ice-cold NLB buffer (10 mL Nuclei EZ lysis buffer supplemented with 1 tablet of protease inhibitor and 10 µl of each RNase inhibitors) and incubated on ice for 5 min with an additional 500 µl of NLB. The tissue and NLB buffer mix was then transferred to the Dounce tissue grinder (D8938, Sigma), gently grind 30 times with Pestle A and 15 times with Pestle B. The homogenate was then incubated on ice for an additional 5 min. The homogenate was filtered through a mini-40-µm and a subsequent mini 20-µm cell strainer (43-10040-50 & 43-10020-50; PluriSelect) and then centrifuged at 500 x *g* for 5 min at 4°C. The pellet was resuspended and washed twice with 1 mL of NLB buffer to remove residue cytoplasmic RNAs. After another centrifugation, the nuclei pellet was resuspended in Nuclei Suspension Buffer (1XPBS, supplemented with 2% BSA and 2 µl Promega RNase inhibitor), filtered through a mini-10-µm cell strainer (43-10010-50, PluriSelect). Nuclei were counted on hemocytometers (KIMA211710, Vetriplast-10).

### Single-nucleus library preparation and sequencing

The quantified brain and muscle nuclei were pooled together in 1:2 ratio, kidney nuclei were processed on its own. The nuclei were partitioned into each droplet with a barcoded gel bead using the 10X Chromium instrument with the Chromium Single Cell 3′ Reagent Kit v.3.1 to generate transcriptome libraries. The libraries were sequenced on the Nova-Seq 6000 (Illumina).

### snGESTALT library preparation

To generate snGESTALT libraries, half of each 10X transcriptome libraries was enriched using in a two-step nested PCR reaction involving:

1) GP6 and READ1 primers (Supplementary Table S5) and Q5 polymerase (NEB), and 2) READ1 and GREAD2 primers (Supplementary Table S5) and Q5 polymerase. The first Q5 reaction (98°C, 30s, 61°C, 25s; 72°C, 30s; 15 cycles) was cleaned up with 0.6X AMPure beads and eluted in 20 µl Ultrapure water (Invitrogen); The second Q5 reaction (98°C, 30s, 61°C, 25s; 72°C, 30s; 8 cycles) was also cleaned up with 0.6X AMPure beads and eluted in 20 µl Ultrapure water (Invitrogen). The concentration of the PCR products was assessed using both High Sensitivity D1000 Screen tape on TapeStation (Aglient) and Qubit™ dsDNA HS Assay Kit. Final PCR was then carried out to incorporate sequencing adapters and sample indexes as described in the Chromium Single Cell 3’Reagent kit v.3.1. The NEBNext Library Quant kit was used for library quantification. Libraries were sequenced using MiSeq (300 cycles kit) and 20% PhiX spike-in. Sequencing parameters: Read1 28 cycles and Read2 300 cycles.

### Processing of raw transcriptome sequencing reads

In brief, sequencing data were demultiplexed and aligned to the zebrafish genome (GRCz10) using Cell Ranger v6.1 with the setting--include-introns. An expression matrix was generated based on UMI counts per gene per cell.

### snRNA-Seq Data Analysis

Seurat v4 ^[38]^ was used for downstream analysis. We analysed each zebrafish separately and removed low quality nuclei with less than 200 genes detected. We also excluded nuclei with a high percentage of UMIs mapped to mitochondrial genes (> 0.02). An R package, SoupX ^[43]^, was used to remove ambient RNA. Briefly, ambient RNA expression is estimated from the empty droplets. The contamination fraction is then calculated with the setting “autoEstCont” followed by the “adjustCounts” function with default parameters to generate a UMI count matrix. We also used scDblFinder^[44]^ to remove doublets. Data were normalised and scaled using SCTransform^[45]^. Mitochondria gene expression was removed from the datasets. We integrated four brain and muscle datasets, and three kidney datasets using integration anchors^[46]^. We then performed dimension reduction, clustering and differential expression analysis on individual clusters using the FindAllMarkers function in Seurat. Canonical marker genes together with the top differentially expressed genes for each cluster were used to identify cell type populations. Pathway analysis was performed using ClusterProfiler^[47]^.

### GESTALT barcode analysis

Sequencing data from GESTALT libraries were processed with the GESTALT pipeline (https://github.com/mckennalab/SingleCellLineage). In brief, read 1 and read 2 were concatenated to generate GESTALT barcode reads with the 10X cell barcodes. A consensus sequence was called by merging all reads that shared the same cell barcode. Consensus sequences were then aligned to a reference sequence using NEEDLEALL aligner with the default settings except`-primersToCheck=FORWARD and-umiLength=28`. The GESTALT barcodes were matched with the corresponding nuclei from the transcriptome datasets based on the 10X cell barcode. Nuclei with <2 GESTALT barcode UMIs were excluded from the lineage trees. Lineage trees were constructed using TreeUtils (https://github.com/mckennalab/TreeUtils) with the setting`--subsetFirstX 4` to anchor the tree with the early edit sites (site 1-4) and extended it with the late edit sites (site 5-9). The trees were then annotated with tissue type.

---

**Extended Data Figure 1.**
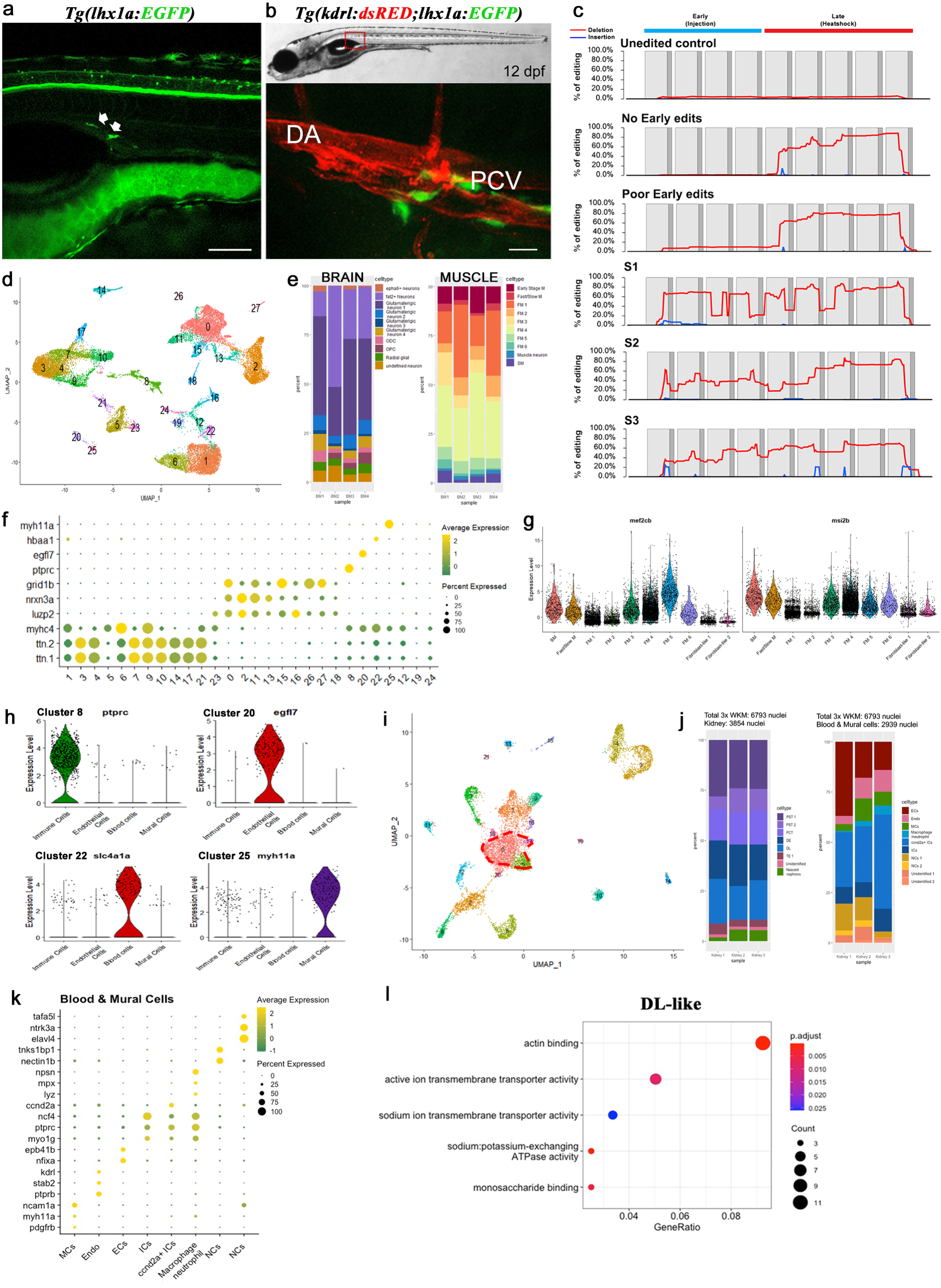
Zebrafish brain, muscle and kidney single nuclei sequencing. **(a)** Side view of a 12 dpf *lhx1a-EGFP* larvae showing EGFP^+^ RPCs around the axial vessels. Scale bar, 100µm **(b)** Close-up of a 12 dpf *Tg(kdrl:dsRed;lhx1a:EGFP)* larvae showing the EGFP^+^ RPCs in close proximity to the vasculature. Scale bar, 20µm **(c)** GESTALT barcode editing efficiency. Editing rate detected in each of the nine CRISPR target sites of the GESTALT barcode plus control. Animals with high editing efficiency at all nine sites were selected for experiments (S1, S2 and S3). **(d)** UMAP showing the initial identification of 31 transcriptionally distinct muscle and brain clusters. **(e)** Percentage of nuclei per cluster from each of the 4 biological replicates of muscle and brain. **(f)** Dotplot showing key markers used to distinguish muscle and brain populations. **(g)** Violin plot showing the expression of *mef2cb* and *msi2b*, indicating the differentiating muscle clusters. **(h)** Violin plot showing the key marker for immune cells, endothelial cells, blood cells and mural cells from both muscle and brain tissue. Each cell type is color coded to match the highlights in UMAP. **(i)** UMAP showing the initial identification of 22 transcriptionally distinct kidney clusters. Cluster 0, 6 and 17 appeared to contain genes from multiple populations, suggesting poor quality that were possibly caused by the heat shock procedure during sample preparation (highlighted in red-dotted circle), thus removed for cluster identification. **(j)** Percentage of nuclei per cluster from each of the 3 biological replicates of kidney. **(k)** Dotplot showing key markers used to identify non-nephron populations. **(l)** Go-term analysis for the DL-like cluster.

**Extended Data Figure 2.**
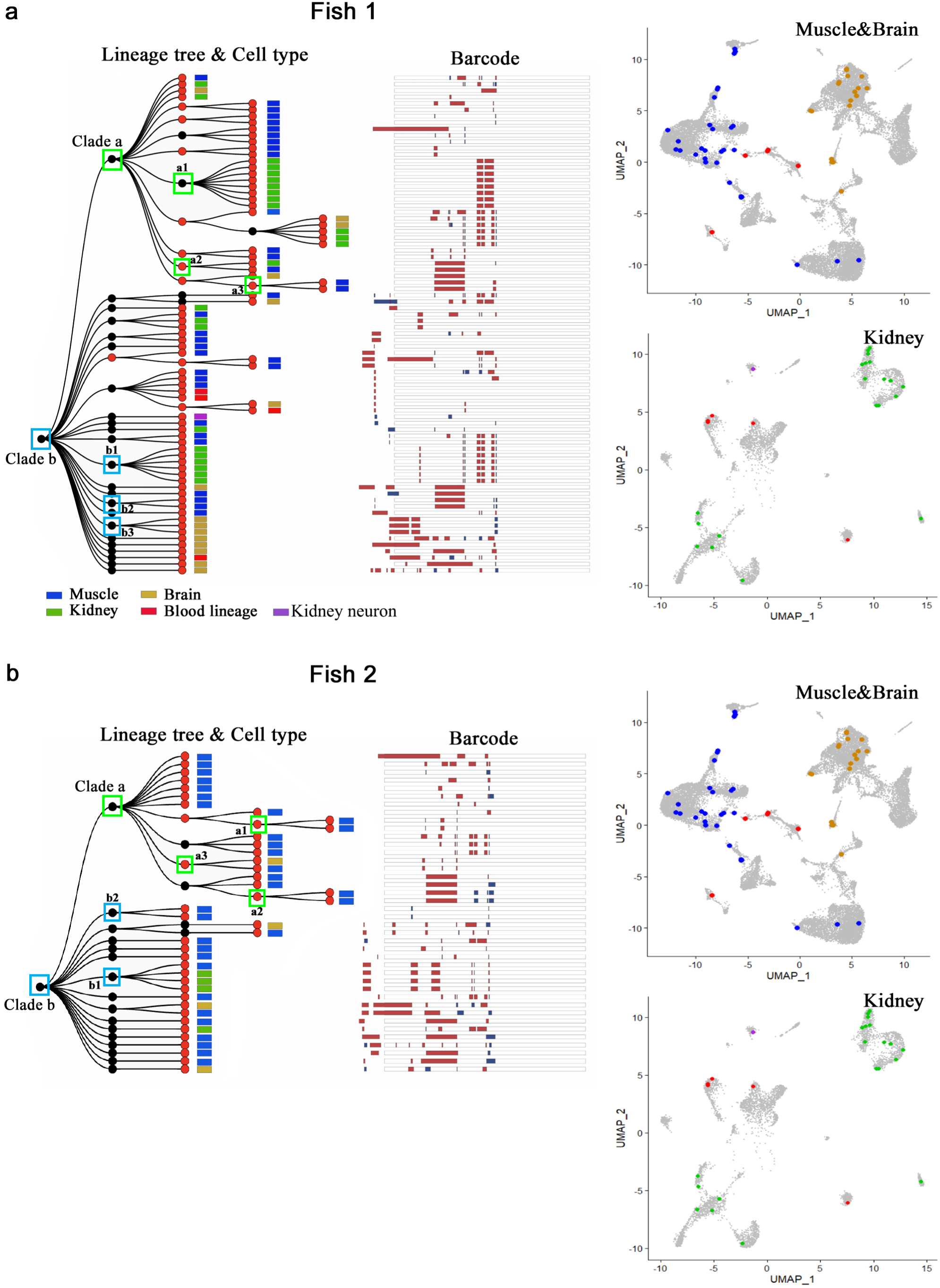
Reconstructed lineage tree of fish 1 and fish 2 kidney, muscle and brain using scGESTALT. **(a)** Barcodes recovered from one juvenile zebrafish kidney, muscle and brain (Fish 1). **(b)** Barcodes recovered from the second juvenile zebrafish kidney, muscle, and brain (Fish 2). The cell lineage trees were assembled based on shared edits using a maximum parsimony approach and threshold set for UMI>2. Black nodes indicate early barcode edits; red nodes indicate late edits. Black lines connect individual cells to nodes on the tree. Cell types (identified from simultaneous transcriptome capture) are color coded as indicated in the legend. The barcode for each cell is displayed as a white bar with deletions (red) and insertions (blue). Green-and blue-colored boxes represent clades ‘a’ and ‘b’, respectively and their subclades, which are shown as mini lineage trees in Figure 2.

**Extended Data Figure 3.**
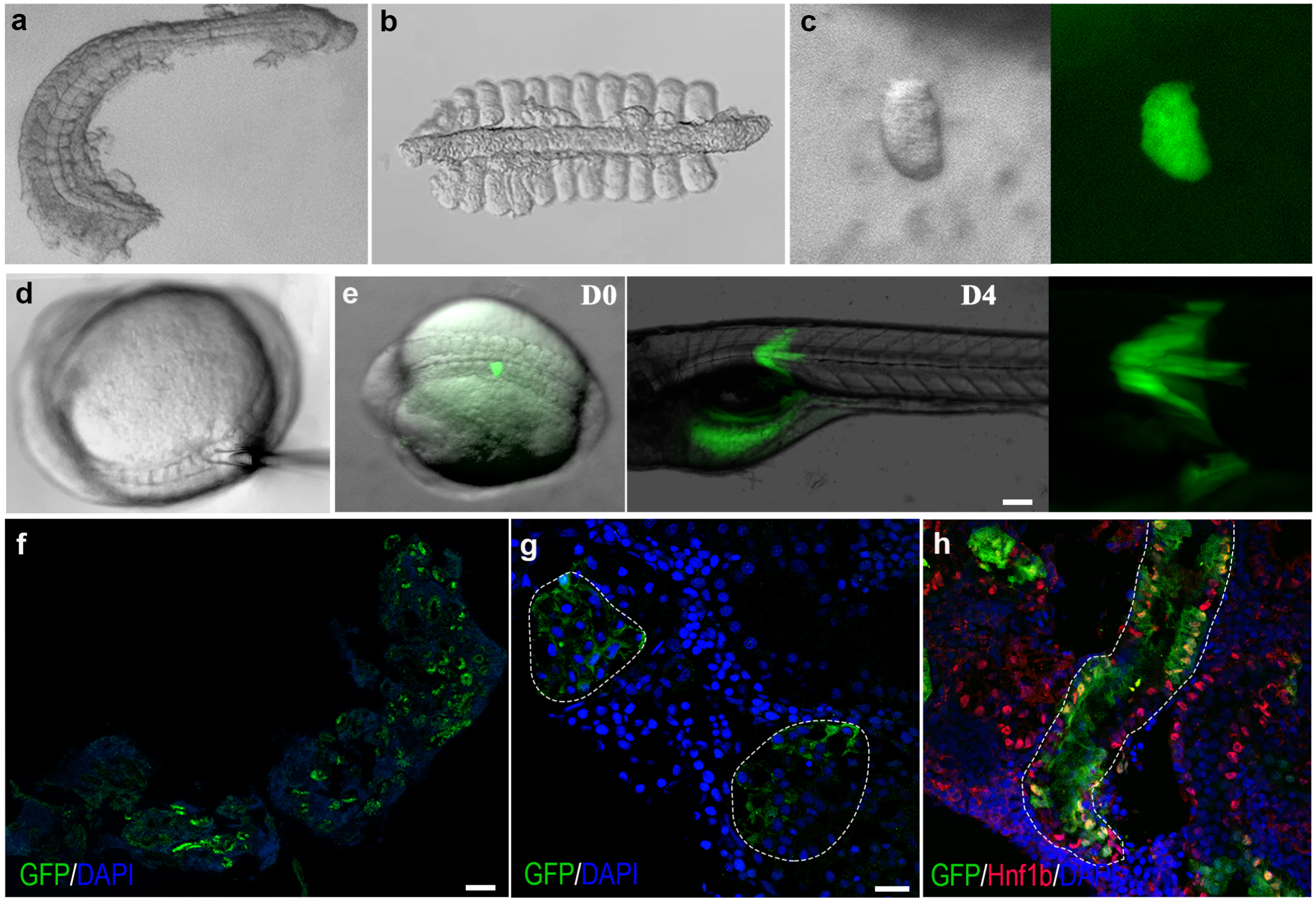
Overview of the somite transplantation procedure and formation of donor-derived nephrons from 18-somite stage. (a-b) Collagenase digested donor embryo. **(c)** An isolated individual somite. **(d)** Image showing the removal of recipient embryo somites using a capillary needle. **(e)** Image showing recipient embryo carrying transplanted donor GFP^+^ somites, and the resulting GFP^+^ muscle fibers at 4 dpf. Scale bar, 100 µm. **(f-g)** Images of transplantation experiment at 18-somites stage, donor somite harvested at 18-somite stage and transplanted into an equivalent stage recipient embryo. **(f)** Low magnification image of a 1.5-month-old recipient fish with GFP^+^ donor derived tubules. Scale bar, 100 µm. **(g)** Donor derived GFP^+^ glomeruli in recipient kidney. Scale bar, 20 µm. **(h)** Donor derived GFP^+^/Hnf1b^+^ tubules in recipient fish. n=2, Scale bar, 20 µm.

**Extended Data Figure 4.**
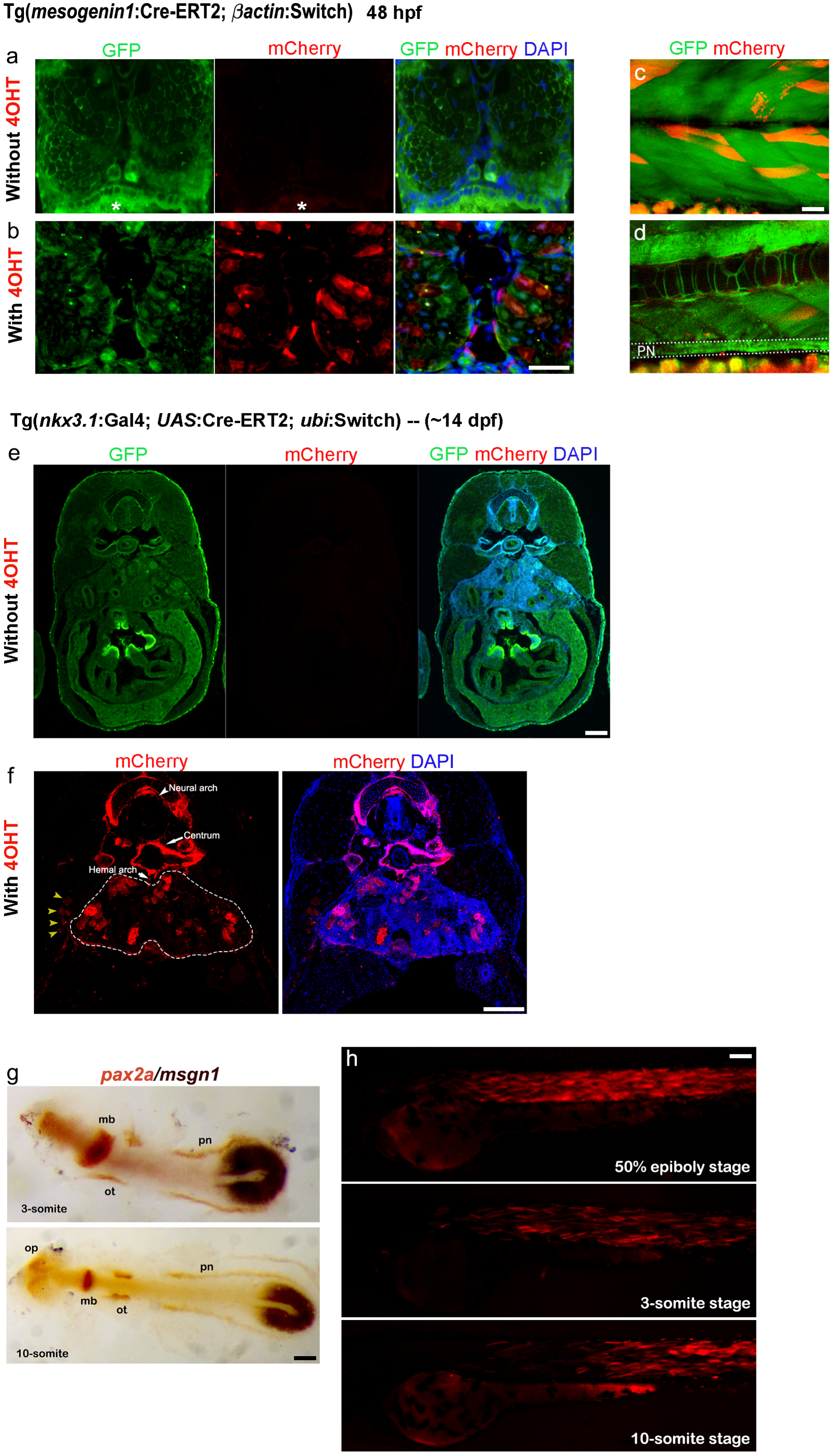
Expression of mCherry in both *msgn1*-Cre and *nkx3.1*-GAL-UAS-Cre fish with and without 4-OHT. (a-b) Cross-section of 7mm *msgn1*-Cre larvae treated without (a) and with (b) 4-OHT. Scale bar, 100 µm. **(c)** Representative confocal image of endogenous mCherry/GFP expression in muscle fibers of live, 4-OHT treated Tg(*msgn1*:CreERT2;*bactin*:Switch) 7mm larvae. **(d)** Image of the same larvae showing lack of mCherry^+^ cells in the pronephros (PN; dotted outline). Scale bar, 20 µm. **(e,f)** Cross-section of 7mm *nkx3.1-GAL-UAS-C*re larvae treated without **(e)** and with **(f**) 4-OHT. Scale bar, 100 µm. **(g)** Double in situ of *pax2a* and *msgn1* of embryos at 3 and 10 somite stage, showing the relative location of intermediate mesoderm (IM, *pax2a*^+^) and paraxial mesoderm (*msgn1*^+^). Scale bar, 100 µm. **(h)** Expression pattern of msgn1 and *pax2a* at 50% epiboly, 3-and 10-somite stages and mCherry expression was not found in pronephros after Cre-loxP recombination. Scale bar, 100 µm.

